## supplemental information for "Remodeling of dermal adipose tissue alleviates cutaneous toxicity induced by anti-EGFR therapy"

**Table S1. Weight loss in EGFRI-related clinical trials.**

| EGFR inhibitors | Trial | Weight loss (%) | |
|---|---|---|---|
| | | All grade | Grade≥3 |
| Afatinib | LUX-Lung 3(Sequist et al., 2013) | 17 | 1 |
| Erlotinib | SATURN("Drug Approval Package: Tarceva (Erlotinib) NDA # 021743," 2010) | 3.9 | < 1 |
| Erlotinib plus gemcitabine | NCIC CTG PA.3("Drug Approval Package: Tarceva (Erlotinib) NDA # 021743," 2010) | 39 | 2 |
| Gefitinib | ARCHER 1050(Wu et al., 2017) | 17 | 0.4 |
| Dacomtinib | ARCHER 1050(Wu et al., 2017) | 26 | 2.2 |
| Neratinib | ExteNET(Mortimer et al., 2019) | 4.8 | 0.1 |
| Neratinib plus capecitabin | NALA(Saura et al., 2020) | 19.8 | 0.3 |
| Lapatinib plus capecitabin | NALA(Saura et al., 2020) | 13.2 | 0.6 |

**Table S2. Blood cell analysis of Vehicle and HFD rats.** WBC: white blood cells. W-SCR: WBC-small cell ratio. W-MCR: WBC-middle cell ratio. W-LCR: WBC-large cell ratio. W-SCC: WBC-small cell count. WBC-MCC: WBC-middle cell count. WBC-LCC: WBC-large cell count. RBC: red blood cells. HGB: Haemohlobin. HCT: Haematocrit. MCV: Mean corpuscular volume. MCH: Mean corpuscular haemoglobin. MCHC: Mean corpuscular haemoglobin concentration. RDW-SD: RBC-distribution width standard deviation. RDW-CV: RBC-distribution width variation coefficient. PLT: Platelets. PDW: Platelets distribution width. MPV: Mean platelet volume. P-LCR: Platelet large cell ratio. Data are presented as the means ±SEM. $P < 0.05$ using 2-tailed unpaired Student's t test.

| | **Ctrl-Afa** | **HFD-Afa** | | **Ctrl-Afa** | **HFD-Afa** |
|---|---|---|---|---|---|
| **WBC** | | | **RBC** | | |
| **WBC (*10^9/L)** | 35.00±7.87 | 20.22±5.23* | **RBC (*10^12/L)** | 9.97±0.08 | 10.35±1.69 |
| **W-SCR** | 0.54±0.05 | 0.57±0.08 | **HGB (g/L)** | 179.00±5.72 | 192.40±34.80 |
| **W-MCR** | 0.14±0.03 | 0.17±0.02 | **HCT** | 0.54±0.02 | 0.56±0.10 |
| **W-LCR** | 0.32±0.02 | 0.26±0.08 | **MCV (fL)** | 53.67±1.28 | 54.52±1.06 |
| **W-SCC (*10^9/L)** | 18.47±2.76 | 11.40±2.75* | **MCH (pg)** | 17.97±0.45 | 18.54±0.75 |
| **W-MCC (*10^9/L)** | 5.10±1.98 | 3.40±0.89 | **MCHC (g/L)** | 334.67±4.03 | 279.80±124.05 |
| **W-LCC (*10^9/L)** | 11.43±3.46 | 5.42±2.57* | **RDW-SD (fL)** | 28.80±1.50 | 27.48±1.61 |
| **PLT** | | | **RDW-CV** | 0.14±0.01 | 0.12±0.01 |
| **PLT (*10^9/L)** | 1373.00±515.77 | 883.40±570.36 | | | |
| **PDW** | 10.75±0.65 | 11.00±0.79 | | | |
| **MPV (fL)** | 9.70±0.10 | 9.40±0.59 | | | |
| **P-LCR** | 0.23±0.01 | 0.24±0.04 | | | |

**Table S3. Blood cell analysis of Vehicle and DCA rats.** WBC: white blood cells. W-SCR: WBC-small cell ratio. W-MCR: WBC-middle cell ratio. W-LCR: WBC-large cell ratio. W-SCC: WBC-small cell count. WBC-MCC: WBC-middle cell count. WBC-LCC: WBC-large cell count. RBC: red blood cells. HGB: Haemohlobin. HCT: Haematocrit. MCV: Mean corpuscular volume. MCH: Mean corpuscular haemoglobin. MCHC: Mean corpuscular haemoglobin concentration. RDW-SD: RBC-distribution width standard deviation. RDW-CV: RBC-distribution width variation coefficient. PLT: Platelets. PDW: Platelets distribution width. MPV: Mean platelet volume. P-LCR: Platelet large cell ratio. Data are presented as the means ±SEM. $P < 0.05$ using 2-tailed unpaired Student's t test.

| | **Vehicle** | **DCA** | | **Vehicle** | **DCA** |
|---|---|---|---|---|---|
| **WBC** | | | **RBC** | | |
| **WBC (*10^9/L)** | 21.92±2.76 | 28.86±1.89** | **RBC (*10^12/L)** | 10.20±1.38 | 7.93±1.01* |
| **W-SCR** | 0.48±0.08 | 0.50±0.06 | **HGB (g/L)** | 198.60±25.45 | 154.40±18.07* |
| **W-MCR** | 0.07±0.03 | 0.08±0.02 | **HCT** | 0.59±0.08 | 0.47±0.05* |
| **W-LCR** | 0.45±0.07 | 0.43±0.05 | **MCV (fL)** | 57.84±1.14 | 58.40±1.77 |
| **W-SCC (*10^9/L)** | 10.52±2.20 | 14.26±1.51* | **MCH (pg)** | 19.50±0.59 | 19.50±0.66 |
| **W-MCC (*10^9/L)** | 1.46±0.63 | 2.24±0.65 | **MCHC (g/L)** | 337.20±4.31 | 334.00±3.03 |
| **W-LCC (*10^9/L)** | 9.94±2.11 | 12.36±1.90 | **RDW-SD (fL)** | 25.86±0.72 | 26.46±0.64 |
| **PLT** | | | **RDW-CV** | 0.11±0.01 | 0.11±0.01 |
| **PLT (*10^9/L)** | 1106.40±576.39 | 1380.60±464.14 | | | |
| **PDW** | 9.34±0.67 | 8.44±0.14* | | | |
| **MPV (fL)** | 8.56±0.43 | 8.12±0.16 | | | |
| **P-LCR** | 0.14±0.03 | 0.09±0.01* | | | |

**Table S4. Primer sequence information**

| Gene | Forward primer | Reverse primer |
|---|---|---|
| **Rat** | | |
| *Adipoq* | GTTCTCTTCACCTACGACCAGTATC | TGGTAGAGAAGGAAGCCTGTAAATG |
| *Plin2* | CTATTCTGAACCAGCCAACATCTGA | TAACTGCTCCTTTGGTCTTATCCAC |
| *Plin4* | GGATACTTCAAATTCTGCATTCCCA | GTCCTCCGTGCCTGTAAGTATATG |
| *Fabp4* | TGACAGGAAAGTGAAGAGCATC | CATGCCCTTTCGTAAACTCTTGTAG |
| *Fads2* | CATCAGCTACTATGCACGTTTCTTC | CATGACAATGTGGTTCATCTGTGTG |
| *Cd36* | AACCACTGAAGAATCTGAAGAGACC | TGAAAGCAACAAACATCACTACTCC |
| *Pparg* | CTTTATGGAGCCCAAGTTTGAGTTT | GTAGCAGGTTGTCTTGAATGTCTTC |
| *Camp* | GATGACTTCAACCAGCAGTCTTTG | CTTGCTACAGACAGTCTCCTTCAC |
| *Col1a1* | GTGGAAACCTGATGTATGCTTGAT | CTTCTGCGTCTGGTGATACATATTC |
| *Col1a2* | GCGATTACTACTGGATTGACCCTAA | CGGCTGTATGCATTCTTGGC |
| *Col5a1* | AGTGTTACCTCCAATTCCTCCAATC | GGTCAAAGTACGGGTCATAGTAGTT |
| *Col5a2* | TGACAAGATAGAGTGCCAAGAAGTG | AAGCCTTTCACCTTTCTTCCTCTAC |
| *Ctgf* | CAAGAGAATACAGGTGCCAGGAA | CAATTTGGACAAGGAGCATCAGC |
| *Atgl* | TCATCATATCGCACTTTAGCTCCAA | ACTGTGATGGTATTCTTCAGCTCAT |
| *Acta2* | CACGCTGAACTATGCTTCTGGA | CCACGCTCAGTCAGGATCTTC |
| *Pdgfra* | GATCGAAGGCAGGCACATTTATATC | TATTGTGCAAGGTTACTTCAGTGTC |
| *Lipe* | AGGGAAATAACAACTATGGAGCCAA | TCATTTGGGAGACTTTGTTTCTGTG |
| *Tgfb1* | ATAGCAACAATTCCTGGCGTTAC | CGTGGAGTACATTATCTTTGCTGTC |
| *Ccl2* | TGGAGAACTACAAGAGAATCACCAG | TCTAATGTACTTCTGGACCCATTCC |
| *Il6* | GTCATTCAGAGCAATACTGAAACCC | CAAGTGCTTTCAAGATGAGTTGGAT |
| *Tnfa* | CATGGATCTCAAAGACAACCAACTG | GCTGACTTTCTCCTGGTATGAAATG |
| **Human** | | |
| *Il6* | AACAACCTGAACCTTCCAAAGATG | GCTTGTTCCTCACTACTCTCAAATC |
| *Tnfa* | TGAGCACTGAAAGCATGATCC | ATCACTCCAAAGTGCAGCAG |
| *Ccl2* | TCATAGCAGCCACCTTCATTCC | GTCTTGAAGATCACAGCTTCTTTGG |

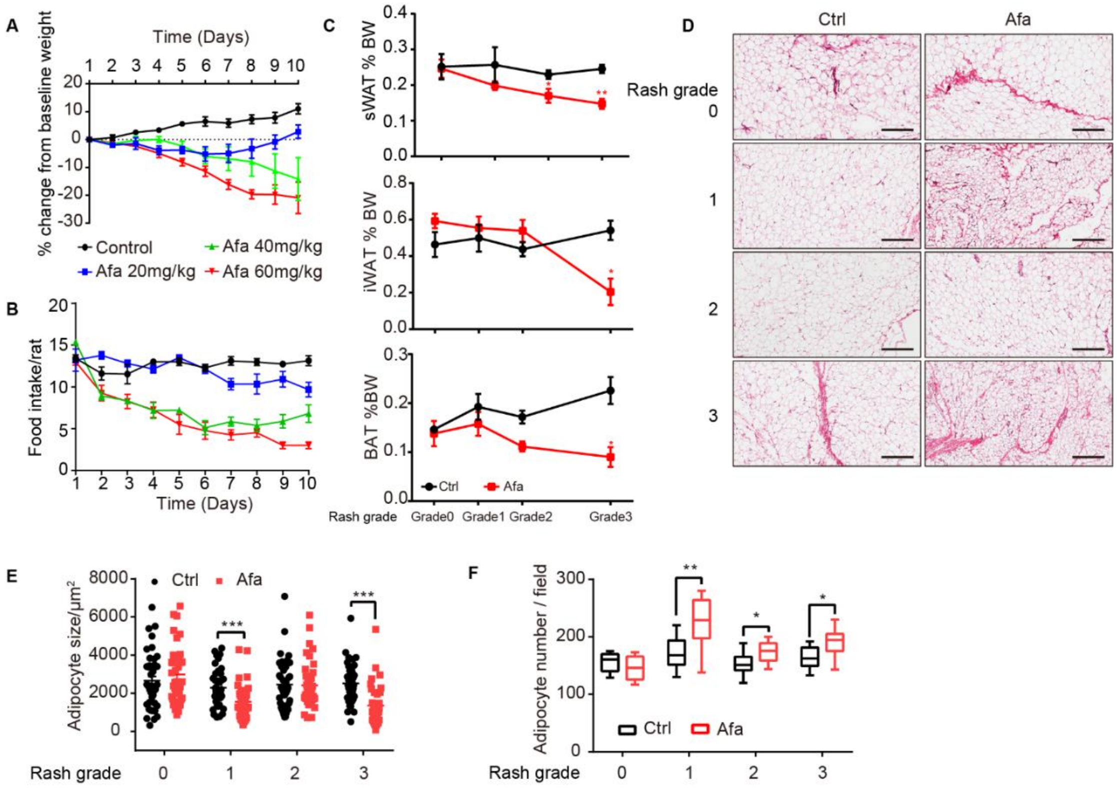


**Figure S1. Characteristics of sWAT during EGFR inhibition**

(A) Body weight change of control and Afa-treated rats. *n*=8 per group. (B) Food intake per rat of different Afa dosage. (C) Percentage of BW change of sWAT, iWAT and BAT. sWAT was obtained under the same shaved-skin area. *n*=3-5 per group. (D) H&E staining of subcutaneous white adipose tissue from Ctrl and Afa-treated rats at indicated time. Scale bars: 300 μm. (E and F) Quantification of the sWAT size (E) and number (F). Data are presented as the means ±SEM. *P < 0.05, **P < 0.01, ***P < 0.001 (Ctrl *vs* Afa) using 2-tailed unpaired Student's t test.

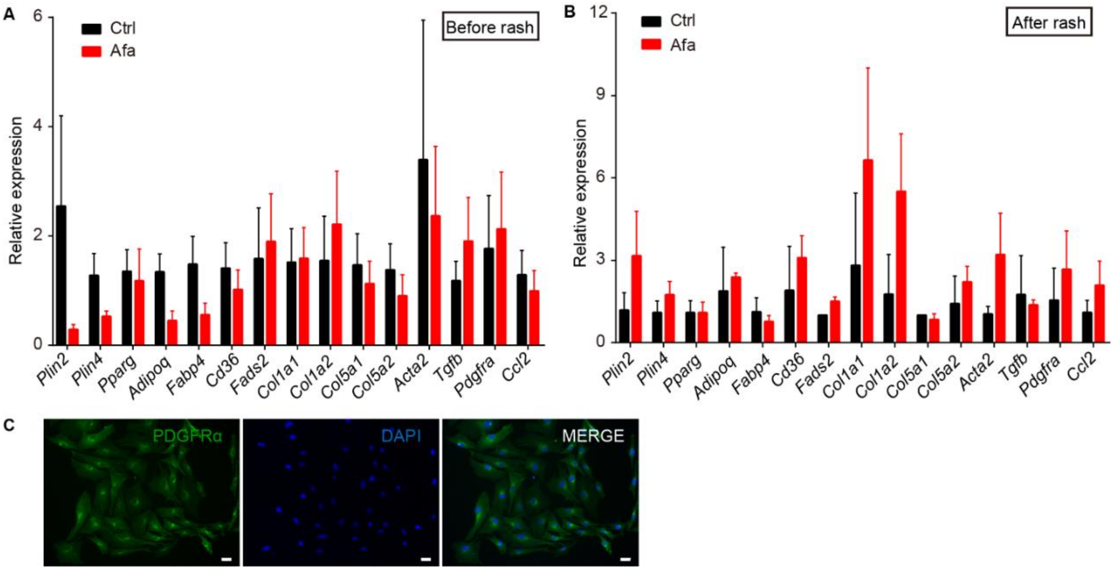


**Figure S2. Additional dedifferentiation changes of dWAT and sWAT**

(A and B) mRNA levels of pro-fibrotic and pro-adipogenic genes in sWAT at one day before (A), and one day after rash (B), *n*=3 per group. (C) PDGFRα (green) and DAPI (bule) immunostaining of dedifferentiated adipocytes under Afa treatment at day 9. Data are presented as the means ±SEM. *P < 0.05, **P < 0.01, ***P < 0.001 (Ctrl *vs* Afa) using 2-tailed unpaired Student's t test.

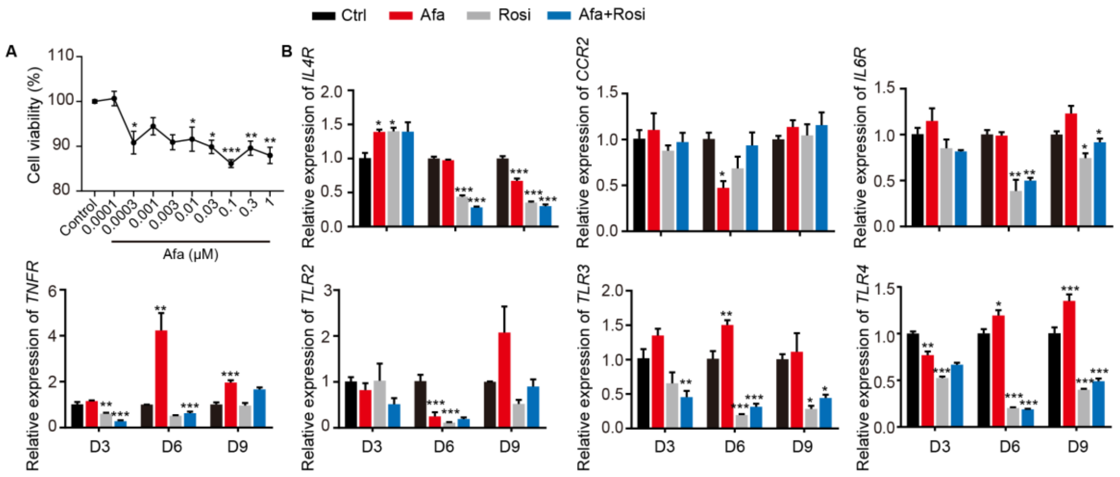


**Figure S3. Expression levels of inflammatory receptors during dFB differentiation**

(A) Viability of dFB cells under different Afa concentrations. (B) mRNA levels of IL4R, CCR2, IL6R, TNFR, and TLR2-3 of differentiating dFB with Afa (10 nM) or Rosi (5 μM) treatment. *n*=4 per group. Data are presented as the means ±SEM. *P < 0.05, **P < 0.01, ***P < 0.001 (Afa/Rosi *vs* Ctrl, Afa+Rosi *vs* Afa) using 2-tailed unpaired Student's t test and one-way ANOVA.

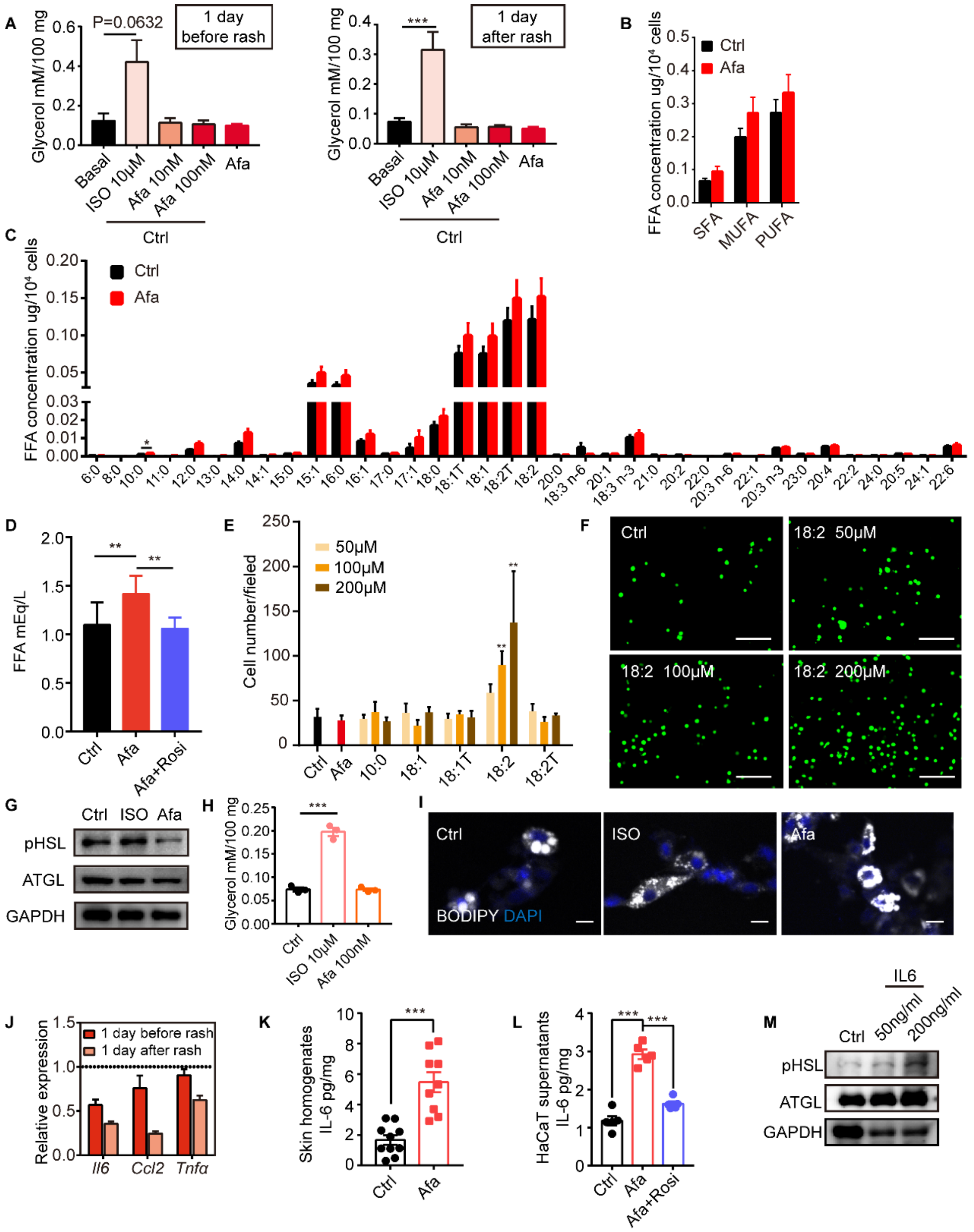


**Figure S4. Lipolytic effects of direct EGFRI stimulation on adipocytes**

(A) Glycerol concentration after 2h ISO or Afa treatment.on isolated sWAT at indicated times. *n*=3-5. (B and C) Lipid mass spectrometry quantification of medium- and long-free fatty acids. *n*=6 per group. SFA: saturated fatty acid . MUFA: monounsaturated fatty acid. PUFA: polyunsaturated fatty acid. (D) FFA concentration of differentiated-dFBs after treatments of HaCaT supernatants from Ctrl, Afa and Afa+Rosi. *n*=3. (E) Quantification of migrated THP-1 cells after stimulation by different FAs. (F) Representative images of migrated THP-1 cells treated with different concentrations of 18:2 FA. Scale bars: 30 μm. (G) Western-blotting of lipase in dFB-derived adipocytes stimulated by 10 μM ISO or 100 nM Afa. (H) Glycerol concentration after 2h ISO or Afa treatment on isolated dWAT. *n*=3. (I) Confocal microscopic images of differentiated dFB cells in basal, ISO and Afa treatment. Lipids were stained with BODIPY 493/503. (J)

Relative mRNA expression of *Il6*, *Ccl2* and *Tnfα* in isolated dWAT. *n*=3 per group. (K) IL6 levels in skin homogenates from Ctrl- and Afa-treated rats. *n*=9-10. (L) IL6 levels in HaCaT supernatants after Ctrl, Afa or Afa+Rosi treatment. *n*=5. (M) Western-blotting of lipase in differentiated-dFB stimulated by Ctrl or IL6. Data are presented as the means. *P < 0.05, **P < 0.01, ***P < 0.001 using 2-tailed unpaired Student's t test.

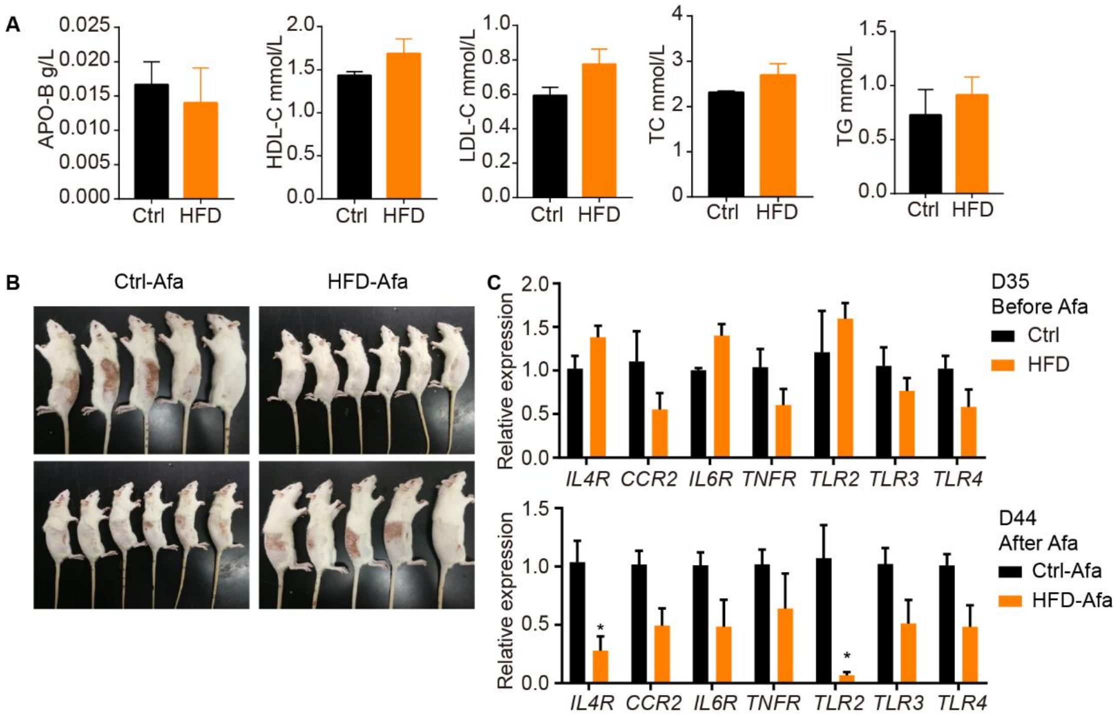


**Figure S5. Additional characterization of HFD rats**

(A) Serum lipid of Ctrl and HFD rats. APO-B refers to apolipoprotein B, HDL-C refers to high-density lipoproteincholesterol, LDL-C refers to low-density lipoproteincholesterol, TC refers to total cholesterol, TG refers to triglyceride. *n*=3 per group. (B) Left and right photos of rash from Ctrl and HFD rats. (C) Relative expression of inflammatory receptors in dFB assayed by RT-PCR from rats at day 35 and day 44. *n*=5 per group. Data are presented as the means ±SEM. *P < 0.05 using 2-tailed unpaired Student's t test.

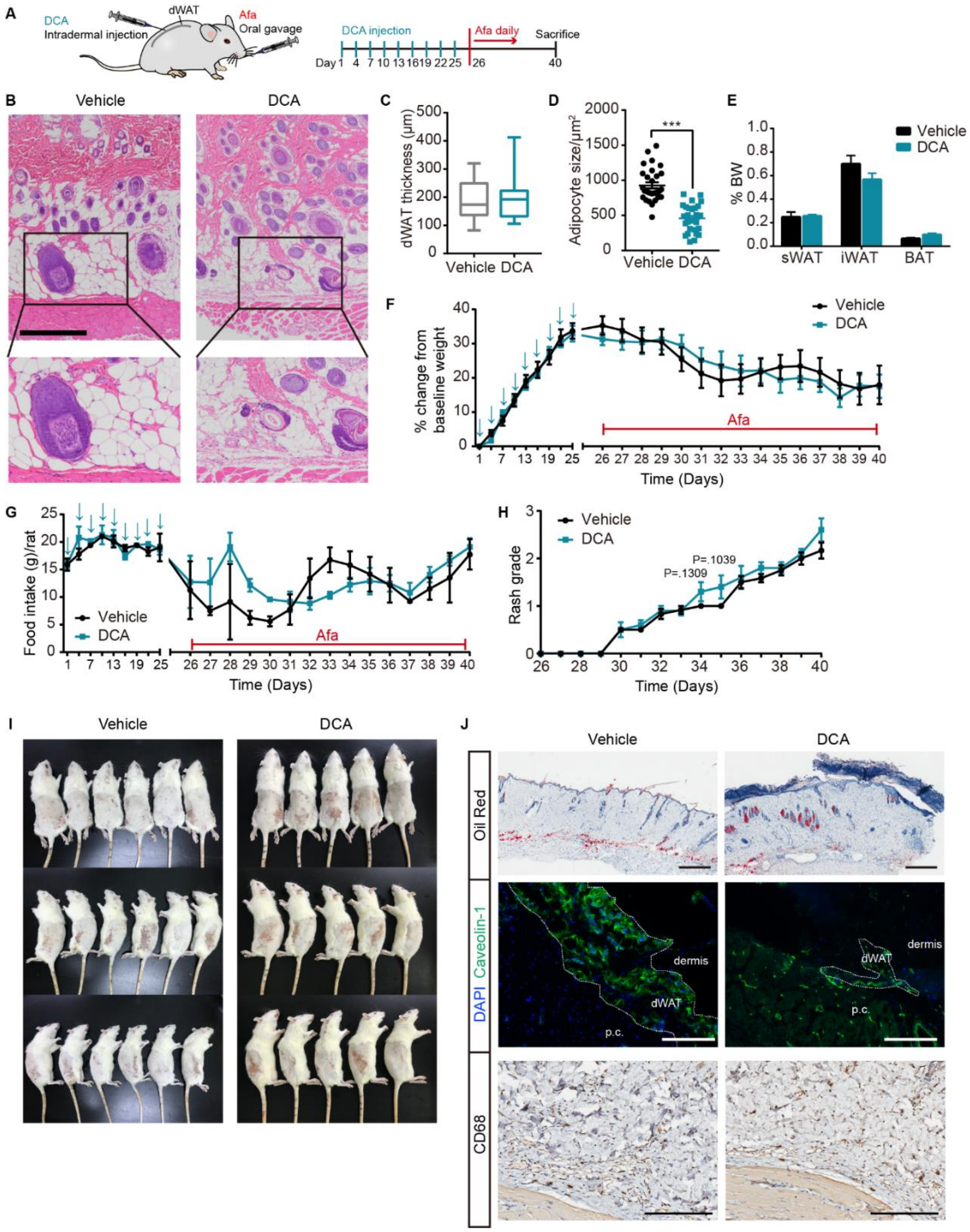


**Figure S6. DCA-induced dWAT ablation aggravates rash phenotypes**

(A) Schematic of the strategy to ablate dWAT by repeatedly DCA intradermal injection. (B) H&E staining of skin from Vehicle and DCA rats. Scale bars: 300 μm. (C and D) Size (C) and thickness (D) of dermal adipocytes. (E) Percentage of BW of sWAT, gWAT and BAT. (F) Body weight. (G) Food intake. (H) Rash grade. (I) Photos of rash at 40 days. (J) Oil Red, Caveolin-1 and CD68 staining of skin biopsies from Ctrl and DCA rats at day 40. Scale bars: 500, 130 and 200 μm (top to down). Data are presented as the means ±SEM. ***P < 0.001 using 2-tailed unpaired Student’s t test.

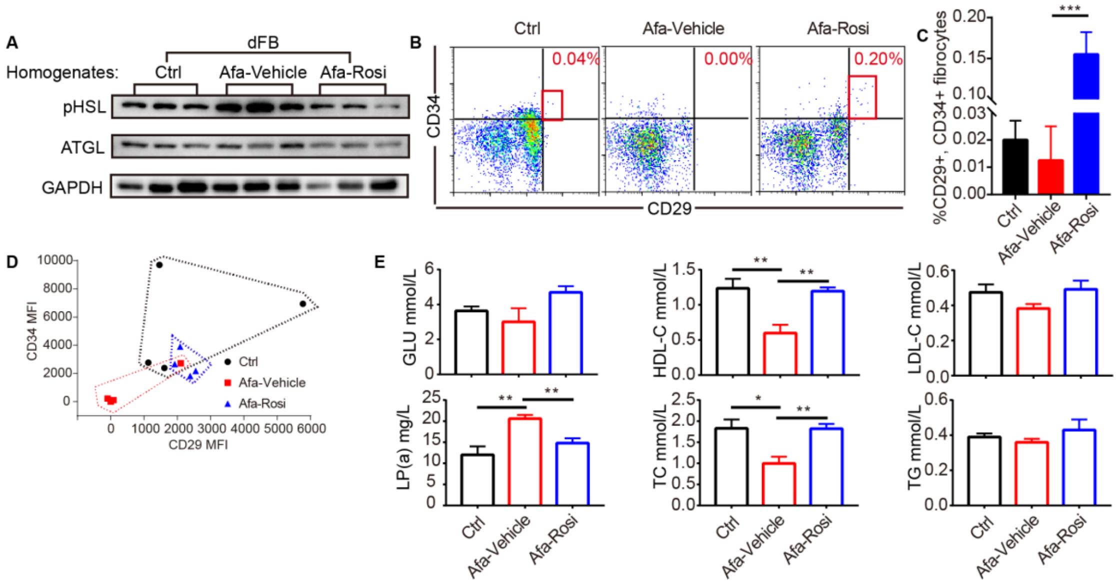


**Figure S7. Additional effects of Rosi prevention in Afa-treated rats**

(A) Western-blotting of lipase in dFB-derived adipocytes stimulated by rat skin homogenates. *n*=3 per group. (B) FACS analysis of APs. (C) Quantification of APs. (D) Mean fluorescence intensity (MFI) of CD29 and CD34. (E) Serum lipid of Ctrl, Afa-Vehicle and Afa-Rosi groups. GLU refers to glucose, HDL-C refers to high-density lipoproteincholesterol, LDL-C refers to low-density lipoproteincholesterol, LP (a) refers to lipoprotein (a), TC refers to total cholesterol, TG refers to triglyceride. *n*=5 per group. Data are presented as the means ±SEM. P < 0.05, **P < 0.01, ***P < 0.001 (Ctrl/Rosi *vs* Afa) using one-way ANOVA.

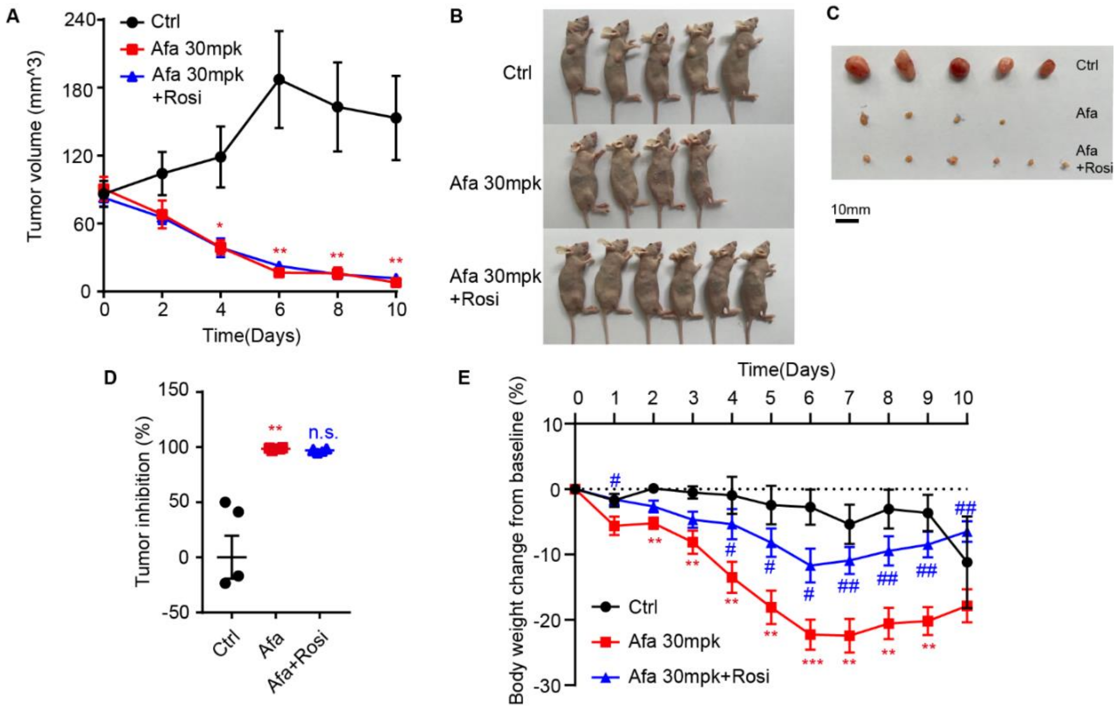


**Figure S8. Rosi did not interfere the anti-tumor effect of EGFRI**

Ctrl: mice gavaged with solvent. Afa 30 mpk (mg/kg): Mice were gavaged with Afa and topically administered vehicle gel. Afa 30 mpk+Rosi: Mice gavaged with Afa and topically administered Rosi gel. (A) Tumor volume. (B and C) Photos of nude mice (B) and tumors (C) at day 10. (D) Tumor inhibition. (E) Body weight. *n*=4-6 per group. Data are presented as the means ±SEM. P < 0.05, **P < 0.01, ***P < 0.001 (Ctrl/Afa+Rosi *vs* Afa) using one-way ANOVA.

**Supplemental Reference**

Drug Approval Package: Tarceva (Erlotinib) NDA # 021743. 2010. *US Food Drug Adm*.

Mortimer J, Di Palma J, Schmid K, Ye Y, Jahanzeb M. 2019. Patterns of occurrence and implications of neratinib-associated diarrhea in patients with HER2-positive breast cancer: analyses from the randomized phase III ExteNET trial. *Breast Cancer Res* **21**:32. doi:10.1186/s13058-019-1112-5

Saura C, Oliveira M, Feng Y-H, Dai M-S, Chen S-W, Hurvitz SA, Kim S-B, Moy B, Delaloge S, Gradishar W, Masuda N, Palacova M, Trudeau ME, Mattson J, Yap YS, Hou M-F, De Laurentiis M, Yeh Y-M, Chang H-T, Yau T, Wildiers H, Haley B, Fagnani D, Lu Y-S, Crown J, Lin J, Takahashi M, Takano T, Yamaguchi M, Fujii T, Yao B, Bebchuk J, Keyvanjah K, Bryce R, Brufsky A, Investigators N. 2020. Neratinib Plus Capecitabine Versus Lapatinib Plus Capecitabine in HER2-Positive Metastatic Breast Cancer Previously Treated With ≥ 2 HER2-Directed Regimens: Phase III NALA Trial. *J Clin Oncol* **38**:3138–3149. doi:10.1200/JCO.20.00147

Sequist L V., Yang JCH, Yamamoto N, O'Byrne K, Hirsh V, Mok T, Geater SL, Orlov S, Tsai CM, Boyer M, Su WC, Bennouna J, Kato T, Gorbunova V, Lee KH, Shah R, Massey D, Zazulina V, Shahidi M, Schuler M. 2013. Phase III study of afatinib or cisplatin plus pemetrexed in patients with metastatic lung adenocarcinoma with EGFR mutations. *J Clin Oncol* **31**:3327–3334. doi:10.1200/JCO.2012.44.2806

Wu Y-L, Cheng Y, Zhou X, Lee KH, Nakagawa K, Niho S, Tsuji F, Linke R, Rosell R, Corral J, Migliorino MR, Pluzanski A, Sbar EI, Wang T, White JL, Nadanaciva S, Sandin R, Mok TS. 2017. Dacomitinib versus gefitinib as first-line treatment for patients with EGFR-mutation-positive non-small-cell lung cancer (ARCHER 1050): a randomised, open-label, phase 3 trial. *Lancet Oncol* **18**:1454–1466. doi:https://doi.org/10.1016/S1470-2045(17)30608-3
