## supplementary file3 for "Remodeling of dermal adipose tissue alleviates cutaneous toxicity induced by anti-EGFR therapy"

**Supplementary file 3. Blood cell analysis of Vehicle and DCA rats.** WBC: white blood cells. W-SCR: WBC-small cell ratio. W-MCR: WBC-middle cell ratio. W-LCR: WBC-large cell ratio. W-SCC: WBC-small cell count. WBC-MCC: WBC-middle cell count. WBC-LCC: WBC-large cell count. RBC: red blood cells. HGB: Haemohlobin. HCT: Haematocrit. MCV: Mean corpuscular volume. MCH: Mean corpuscular haemoglobin. MCHC: Mean corpuscular haemoglobin concentration. RDW-SD: RBC-distribution width standard deviation. RDW-CV: RBC-distribution width variation coefficient. PLT: Platelets. PDW: Platelets distribution width. MPV: Mean platelet volume. P-LCR: Platelet large cell ratio. Data are presented as the means ±SEM. P < 0.05 using 2-tailed unpaired Student’s t test.

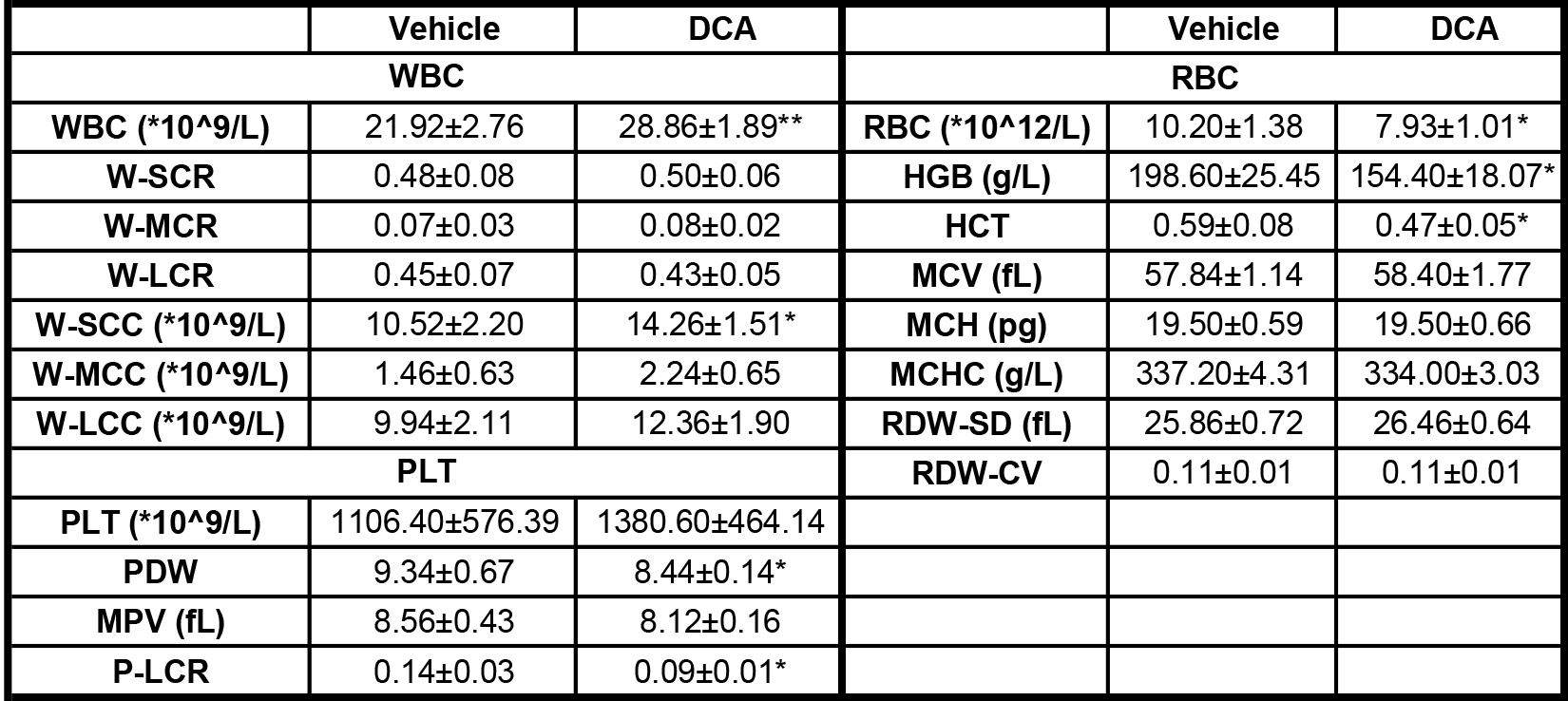
